# Maternal RNA-directed DNA methylation is required for seed development in *Brassica rapa*

**DOI:** 10.1101/174805

**Authors:** Jeffrey W. Grover, Timmy Kendall, Abdul Baten, Graham J. King, Rebecca A. Mosher

**Affiliations:** Department of Molecular & Cellular Biology, The University of Arizona, Tucson, AZ 85721, USA; The School of Plant Sciences, The University of Arizona, Tucson, AZ 85721, USA; Southern Cross Plant Science, Southern Cross University, Lismore, NSW 2480, Australia

**Keywords:** RNA-directed DNA Methylation, Polymerase IV, Polymerase V, siRNA, seed development

## Abstract

Some organisms deploy small RNAs from accessory cells to maintain genome integrity in the zygote, a mechanism that has been proposed but not demonstrated in plants. Here we show that maternal mutations in the Pol IV-dependent small RNA pathway cause abortion of developing seeds in *Brassica rapa*. Surprisingly, small RNA production is required in maternal somatic tissues, but not in maternal gametes or the developing zygote. We propose that parental influence over zygotic genomes is a common strategy in eukaryotes and that outbreeding species such as *B. rapa* are key to understanding the role of small RNAs during reproduction.

---

Small RNAs are a common mechanism to protect a genome, with most eukaryotes deploying at least one variety of small RNA to transcriptionally or post-transcriptionally silence transposons (Malone and Hannon 2009; Holoch and Moazed 2015). Suppression of transposons is particularly important in the germ line to prevent inheritance of reactivated elements or transposon-based mutations (Iwasaki et al. 2015). In Drosophila, Piwi-associated (pi)RNAs are produced in pre-meiotic nurse cells that surround and support the egg cell and are loaded into the egg before fertilization (Brennecke et al. 2008; Klattenhoff and Theurkauf 2008). These piRNAs trigger additional piRNA production after fertilization to control transposons in the zygotic germ line. The maternal initiation of piRNA-mediated silencing means that transposons inherited solely from the paternal genome can evade silencing and retrotranspose in the zygote.

Plants do not encode piRNAs, and instead use short interfering (si)RNAs to silence transposons (Matzke and Mosher 2014; Fultz et al. 2015). In the Arabidopsis male gametophyte, 21 nucleotide (nt) siRNAs are produced from transposon sequences in the pollen vegetative nucleus, a haploid cell that supports the sperm cells but does not contribute to the zygote (Slotkin et al. 2009). These siRNAs are transferred to the sperm cells, where they silence homologous transcripts post-transcriptionally (Martínez et al. 2016). However, it is unclear whether pollen siRNAs have any function in the resulting zygote since there are no phenotypes associated with loss of these siRNAs in Arabidopsis.

On the maternal side, 24 nt Pol IV-dependent (p4-)siRNAs are produced from transposons by the sequential action of RNA Polymerase IV, RNA-dependent RNA polymerase 2 (RDR2), and DCL3. P4-siRNAs then associate with ARGONAUTE4 and RNA Polymerase V and transcriptionally silence transposons in a process called RNA-directed DNA methylation (RdDM) (Matzke and Mosher 2014). P4-siRNAs are expressed highly from the maternal genome during Arabidopsis seed development (Mosher et al. 2009; Pignatta et al. 2014) and have been implicated in controlling the effective parental dosage during seed development (Lu et al. 2012; Autran et al. 2011; Erdmann et al. 2017), and in establishing boundaries between euchromatin and heterochromatin (Zemach et al. 2013; Li et al. 2015). However, it is not clear whether p4-siRNAs are produced from the female gametophyte, the filial tissues (embryo and endosperm), or the somatic tissues surrounding these reproductive structures (Mosher et al. 2009).

Neither paternal 21 nt siRNAs nor maternal p4-siRNAs are required for seed development in Arabidopsis (Mosher et al. 2009; Martínez et al. 2016). However Arabidopsis is an inbreeding species with few active transposons and limited parental conflict and therefore might be a poor system to study the function of small RNAs during reproduction. Pol IV and Pol V mutants in tomato display pleiotropic phenotypes, including reproductive defects (Gouil and Baulcombe 2016), suggesting that p4-siRNAs and RdDM might be important during sexual reproduction in genomes with higher transposon content. Here we show that loss of p4-siRNAs in *Brassica rapa* causes a specific defect during seed production. P4-siRNA production is required in maternal somatic tissue for development of embryo and endosperm, suggesting communication between diploid generations in the seed.

## Results and Discussion

To better understand the role of p4-siRNAs during reproduction, we isolated mutations impacting the production and activity of p4-siRNAs in *B. rapa*, an outbreeding relative of Arabidopsis whose genome is approximately 40% transposons (Wang et al. 2011). In Arabidopsis, NRPD1 (the largest subunit of Pol IV) and RDR2 are required for p4-siRNA production, while NRPE1 (the largest subunit of Pol V) produces nascent transcripts that are required for p4-siRNA activity (Matzke and Mosher 2014). Each of these genes has reverted to a single copy in the mesopolyploid *B. rapa* genome (Huang et al. 2013) and we identified putative loss-of-function mutations in each gene (Supplemental Table 1).

After repeated backcrossing to remove other mutations, we sequenced small RNA transcriptomes from the mutants (Supplemental Table 2). As in other plants, 24 nt siRNAs are the most abundant size class in *B. rapa* reproductive tissues (Figure 1A). These siRNAs are strongly reduced in *braA.nrpd1.a-2* and *braA.rdr2.a-2* (hereafter *braA.nrpd1* and *braA.rdr2*), indicating they are p4-siRNAs. Compared to the almost complete loss of 24 nt siRNAs in *braA.rdr2*, *braA.nrpd1* retains a low level of 24 nt siRNAs, suggesting that this allele might not be null mutation. As in Arabidopsis (Mosher et al. 2008; Wang and Axtell 2017), *braA.nrpe1.a-1* reduces, but does not eliminate, *B. rapa* 24 nt siRNAs. As expected, 87% of siRNA-generating loci require *BraA.RDR2* (Figure 1B), indicating that p4-siRNA loci are the predominant class of small RNA-generating loci in *B. rapa*. Many of these loci are strongly downregulated in *braA.nrpe1* (Figure 1B), demonstrating that although this mutation has a small effect on total p4-siRNA accumulation, it has a strong effect on siRNA production at specific loci.

**Figure 1.**
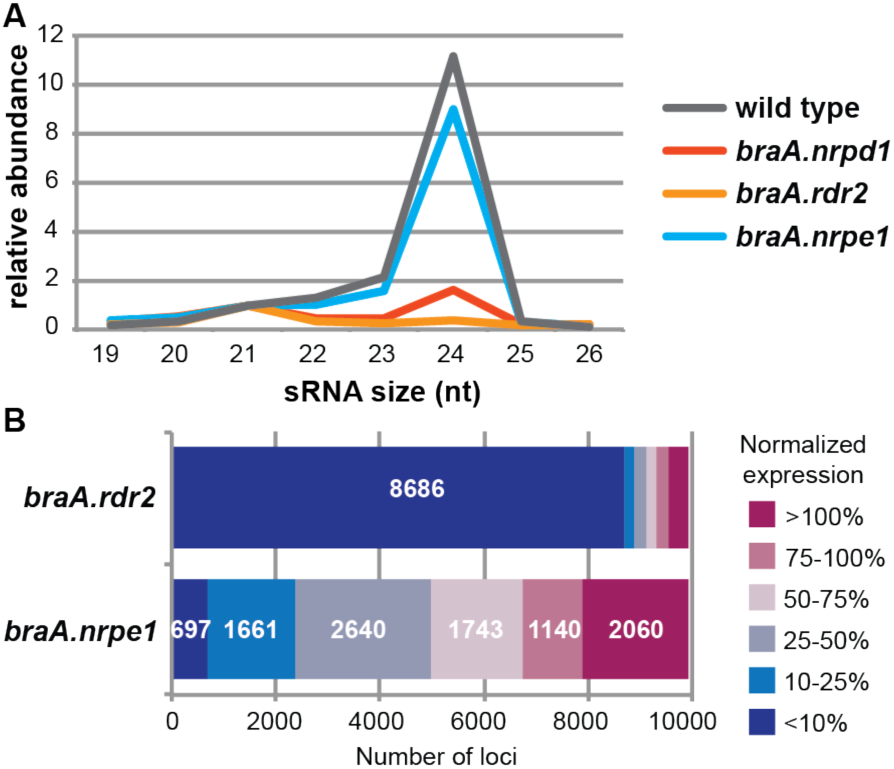
Small RNA analysis of *B. rapa* RdDM mutants. (A) Size profile of genome-matching reads demonstrates that 24 nt small RNAs are the most abundant class of small RNAs in unfertilized ovules. These sequences are lost or strongly reduced in the *braA.rdr2* and *braA.nrpd1* mutants, indicating they are p4-siRNAs. P4-siRNAs are slightly reduced in *nrpe1* ovules. (B) The majority of small RNA-generating loci have sharply reduced expression in *braA.rdr2*, while there is a range of reduced expression in the *braA.nrpe1* mutant.

We observed no growth defects associated with loss of RdDM in *B. rapa*, although *braA.nrpe1* plants were somewhat late flowering. All three mutants developed normal flowers and their siliques (seed pods) enlarged following self-fertilization. Mutant siliques, however, remained slightly smaller than wild-type siliques and produced many fewer viable seeds (Figure 2A). The few seeds that were produced from homozygous mutant plants were smaller, lighter, and less uniformly round when compared to wild type seeds (Figure 2BC). However, these seeds were viable and produced plants that were indistinguishable from first generation homozygous mutants. Hence, mutant seeds that progressed through embryogenesis went on to have normal vegetative development. These second-generation homozygous mutants produced viable seed at a similar frequency to their first-generation parents, suggesting that there is neither a progressive defect associated with loss of p4-siRNAs nor a compensatory change in seeds produced from p4-siRNA mutants.

**Figure 2.**
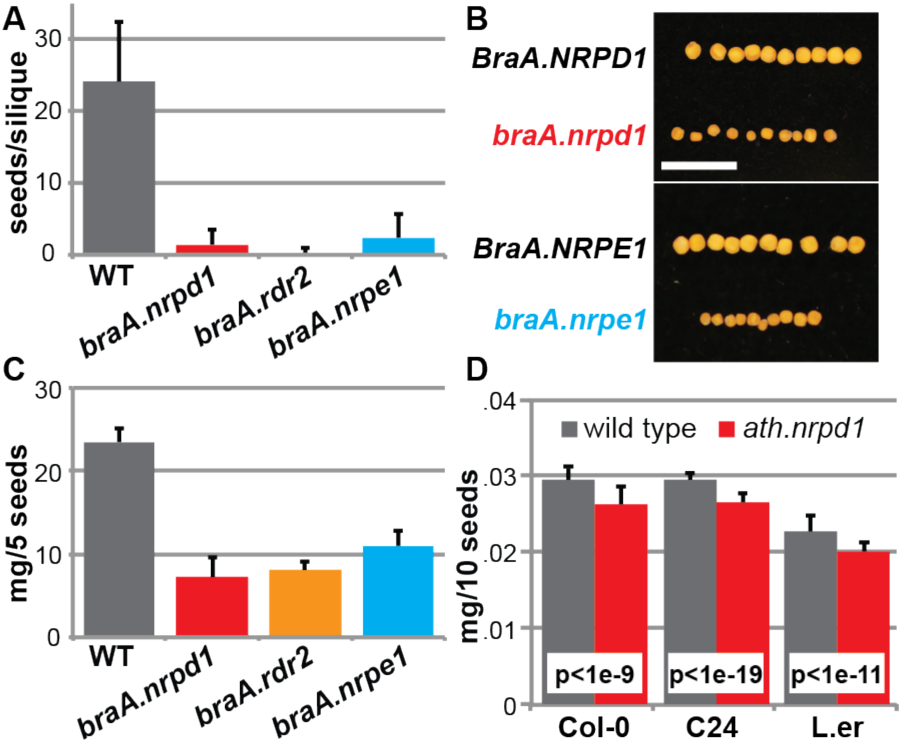
RdDM mutants are defective for seed production. (A) The number of seeds per silique is strongly reduced in all three *B. rapa* RdDM mutants (n > 200 siliques from at least two individuals). (B) Seed from *B. rapa* RdDM mutants are smaller and less uniform in shape than seed from wild-type siblings (scale bar = 10 mm). (C) *B. rapa* mutant seeds are also significantly lighter than wild type (n > 24 measurements). (D) Arabidopsis *nrpd1* mutant seeds are significantly lighter than wild type in three different genetic backgrounds (n > 35 measurements). All error bars depict standard deviation.

Careful assessment of seed weight in Arabidopsis *braA.nrpd1* mutants uncovered a small but significant reduction in seed weight in each of three different genetic backgrounds (Figure 2D). This pattern indicates that p4-siRNAs might have a conserved influence on seed production that varies only in severity between *B. rapa* and Arabidopsis.

To better understand what caused the reduced seed production in *B. rapa* p4-siRNA mutants, we observed early events during seed development. Upon self-fertilization or manual pollination, mutant siliques elongated rapidly and remained indistinguishable from wild type for at least two weeks. Light microscopy of cleared ovules confirmed that fertilization occurred and that the embryogenesis program began correctly (Figure 3B). By day 14 (torpedo stage of embryogenesis), many of the developing mutant seeds were noticeably smaller than wild-type seeds, and smaller and/or shriveled ovules became more abundant as development progressed (Figure 3A). By 17 days after fertilization, diverse seed phenotypes were observed in mutant siliques, ranging from clearly aborted to nearly normal (Figure 3C). However, even the roundest and plumpest mutant seeds were smaller than age-matched wild-type seeds and contained embryos of diverse sizes and developmental states (Figure 3D). Endosperm from mutant seeds was also variable in size and developmental stage. This variability remained evident in mature mutant siliques, which contained a range of aborted seed – from small flecks of completely degenerated seeds to large, but clearly shriveled and unviable seeds. This diversity suggests that in *B. rapa*, loss of p4-siRNAs or their activity causes asynchronous seed abortion, rather than a defect at one fixed point of development.

**Figure 3.**
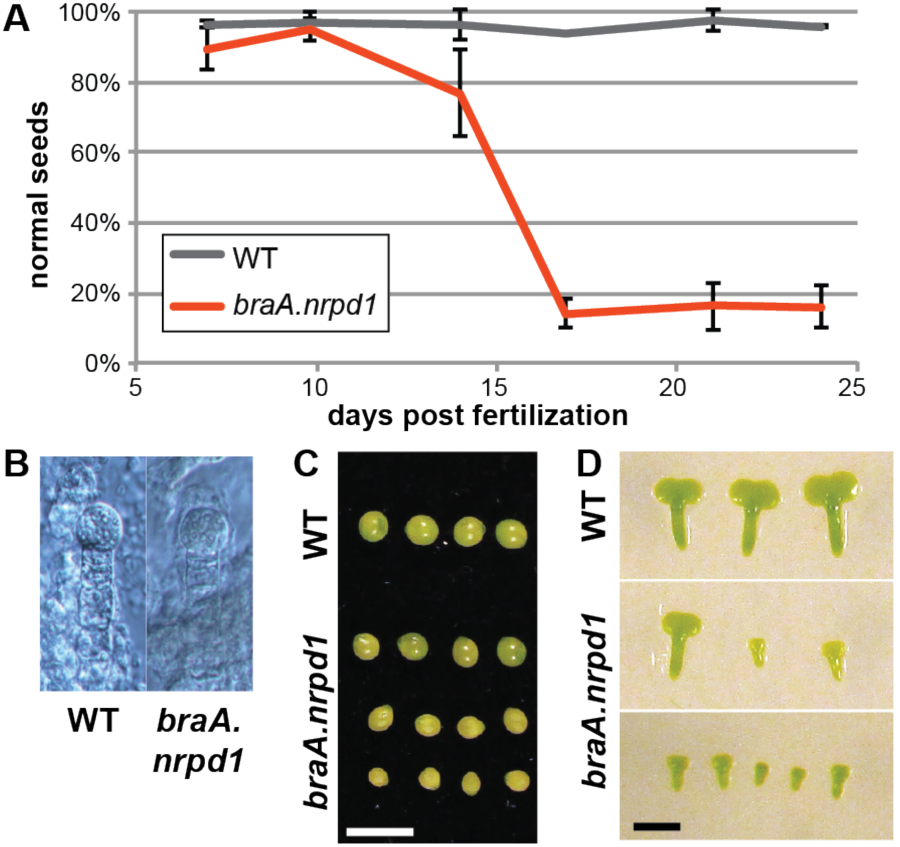
RdDM mutants display asynchronous seed abortion. (A) Siliques were dissected at 7-24 days post fertilization (dpf) and the development of seeds was assessed. Seeds were considered abnormal if they were substantially smaller than wild type or beginning to shrivel. 50-300 seeds were counted for each of 2-3 replicates for each data point. Error bars depict standard deviation of biological replicates. (B) Wild-type and mutant seeds contain identical globular embryos at 7 dpf. (C) Wild-type seeds are uniformly green, round, and plump, while *braA.nrpd1* seeds are variable in size, color, and turgidity at 17 dpf. Even the largest *braA.nrpd1* seeds (top row) are smaller than wild-type seeds and contain smaller and developmentally retarded embryos. The lower two rows of *braA.nrpd1* seeds demonstrates the variation in seed size. Scale bar = 5 mm. (D) Embryos dissected from 17 dpf wild-type or wild-type-like *braA.nrpd1* seeds. Wild-type embryos are uniform in size and developmental stage (late torpedo) while *braA.nrpd1* embryos are variable. Scale bar = 1 mm.

In Arabidopsis, as in Drosophila, the maternal genotype determines whether small RNAs are produced after fertilization; developing seeds with *nrpd1* mothers have strongly reduced p4-siRNA expression even when fertilized by wild-type pollen (Mosher et al. 2009; Lu et al. 2012). To determine whether the reproductive defect in *B. rapa* is controlled by maternal or zygotic genotype, we reciprocally crossed homozygous mutants to wild type and scored seed production. The genotype of pollen donors had no impact on the seed set, while seed production in a mutant x wild type cross (female genotype listed first in all crosses) was indistinguishable from a mutant x mutant cross (Figure 4A). Furthermore, the seeds produced from mutant x wild type crosses were smaller and less round than seeds produced from the reciprocal wild type x mutant cross, indicating that the maternal genotype is responsible for both aspects of the reproductive defect – seed number and seed quality.

**Figure 4.**
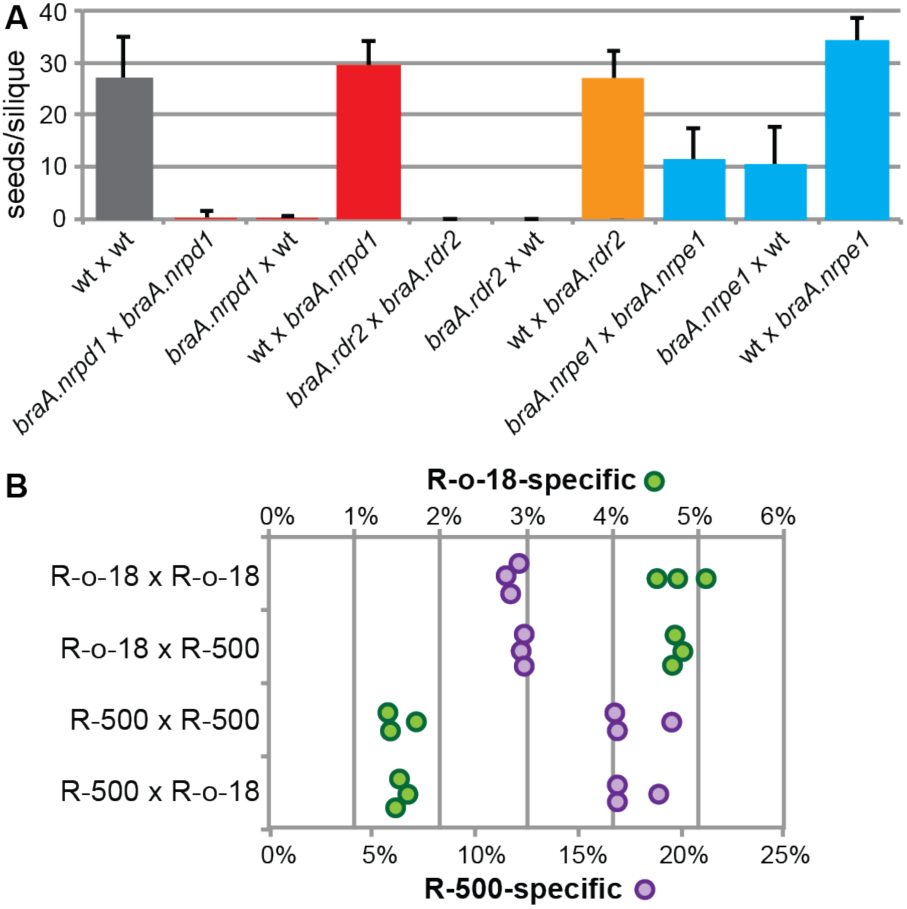
Seed production and siRNA populations are maternally controlled in *B. rapa*. (A) Reciprocal crosses between RdDM mutants and wild type demonstrates that maternal genotype is the determining factor in seed production. Viable seeds were counted for < 10 siliques per cross (maternal parent listed first). Error bars are standard deviation. (B) Seeds from reciprocal R-o-18 x R-500 crosses were dissected 14 days after fertilization and subjected to small RNA sequencing. The resulting libraries (three biological replicates per genotype) were mapped onto both parental genomes. Reads matching only to the R-o-18 genome (green) and those matching only the R-500 (purple) demonstrate that paternal genotype does not substantially influence the small RNA population.

Although in Arabidopsis dissected embryo and endosperm show little parental bias (Pignatta et al. 2014), developing seeds contain overwhelmingly maternal small RNA populations (Mosher et al. 2009). To assess parental contribution of small RNAs in *B. rapa*, we sequenced small RNAs from reciprocal intraspecific (R-o-18 x R-500) crosses at 14 days after fertilization and mapped small RNAs to both genomes independently. Although genome-specific siRNAs do not provide a strict measure of genome-of-origin because of missing sequence in each assembly, comparing the relative level of genome-specific reads between crosses gives an indication of the contribution from the maternal and paternal genomes. As in Arabidopsis, genome-specific reads do not change substantially with different paternal genomes, indicating that the paternal chromosomes do not contribute significantly to the small RNA transcriptome of the seed as a whole (Figure 4B).

We previously hypothesized that p4-siRNAs are maternal-specific after fertilization due to epigenetic patterns established by p4-siRNAs in the female gametophyte before fertilization (Mosher et al. 2009; Mosher 2010). However, perturbation of various epigenetic marks did not influence maternal-specific p4-siRNA production (Mosher et al. 2009; 2011), raising the possibility that p4-siRNAs in the developing seed are instead produced from maternal sporophytic tissue. The clear and quantitative phenotype associated with loss of p4-siRNAs in *B. rapa* provides an excellent system to determine whether p4-siRNAs are required in the diploid maternal sporophyte, the haploid maternal gametophyte, or both. To address this question, we genotyped progeny from crosses with heterozygous mutants, which have a wild-type allele in the sporophyte and in 50% of the resulting gametophytes. A gametophytic defect would result in partial loss of seed production and distortion of the expected allelic ratio in the progeny. Instead, we observed normal seed development and detected the expected 1:1 ratio of alleles in all crosses, indicating that both male and female gametophytes function normally even when carrying a p4-siRNA mutation (Table 1). Combined with the observation that homozygous mutant mothers have defective seed production, we conclude that the seed production defect is due to loss of p4-siRNA production or activity in the maternal sporophytic tissue, even though the defect manifests after successful gametogenesis and fertilization.

**Table 1.** Seed abortion is controlled by maternal sporophytic genotype.

|  | progeny genotype<sup>1</sup> |  |  | p-value<sup>2</sup> |
| --- | --- | --- | --- | --- |
|  | +/+ | +/- | -/- |  |
| Het. x wild type (WT) |  |  |  |  |
| *nrpd1-2/+* x WT | 32 | 34 |  | n.s. |
| *nrpe1-1/+* x WT | 25 | 31 |  | n.s. |
| *rdr2-2/+* x WT | 30 | 20 |  | n.s. |
| wild type (WT) x het. |  |  |  |  |
| WT x *nrpd1-2/+* | 29 | 34 |  | n.s. |
| WT x *nrpe1-1/+* | 35 | 31 |  | n.s. |
| WT x *rdr2-2/+* | 25 | 30 |  | n.s. |
| Het. x homo. |  |  |  |  |
| *nrpd1-2/+* x *nrpd1-2* |  | 68 | 57 | n.s. |
| *nrpe1-1/+* x *nrpe1-1* |  | 65 | 60 | n.s. |
| *rdr2-2/+* x *rdr2-2* |  | 65 | 57 | n.s. |

<sup>1</sup> +/+, WT; +/-, heterozygous; -/-, homozygous mutant
<sup>2</sup> P-values from chi-squared test. n.s., not significant

The requirement for functional p4-siRNA in maternal sporophytic tissues does not rule out the possibility that p4-siRNAs are also required in zygotic tissues. To measure any defect in seed production due to the zygotic genotype we measured seed production and genotypes in crosses between heterozygous mothers and homozygous mutant fathers. Seed production was normal and resulted in equal development of heterozygous and homozygous mutant progeny, confirming that seed abortion is controlled solely by maternal sporophytic genotype (Table 1).

Our analysis of *B. rapa* mutants demonstrates that p4-siRNA production or activity is required in *B. rapa* maternal sporophytic tissue for successful seed development. This is reminiscent of the maternal sporophytic requirement for *NRPD2* (the second subunit of Pol IV and V) for suppression of maternally-transmitted *EVADE* retroelements before meiosis in Arabidopsis (Reinders et al. 2013). While it is possible that seed abortion in *B. rapa* RdDM mutants is due to transposition of reactivated elements before meiosis or gametogenesis, we think this hypothesis is unlikely to be correct because we observe normal gametogenesis, fertilization, and early embryogenesis in RdDM mutant mothers (Figure 3B). It is also possible that RdDM is required specifically during integument development, however it is not clear why epigenetic control of transposons would be important specifically in this tissue.

Instead, we propose that p4-siRNAs produced from maternal sporophytic tissue might move to the developing endosperm and/or embryo to control transposons during seed development. Massive deregulation of transposons would explain the asynchronous and nearly complete seed abortion we observed. It is also possible that mobile maternal p4-siRNAs are required for correct expression of specific transposon-associated genes, such as Agamous-like transcription factors, which are upregulated in Arabidopsis seeds lacking maternal *NRPD1* (Lu et al. 2012). Finally, mobile maternal p4-siRNAs might function across the genome to maintain the effective parental dosage in endosperm and their loss could unbalance parental contributions and lead to endosperm failure (Gehring 2013; Yang et al. 2017).

Our data suggest that control of developing progeny by parental somatic tissues is a more common mechanism than was previously appreciated. In addition to maternal loading of piRNAs in flies, ciliates also use small RNAs from the parental macronuclei to program the macronuclei of daughter cells. Our results offer another example of parental tissues influencing developing progeny through small RNAs, and suggest this is a common biological mechanism.

## Materials & Methods

### Plant growth, genotyping, and phenotyping

All plants were grown in a greenhouse at 18°C with supplemental lighting to 16 hours daylight. The TILLING population was created in R-o-18, an inbred line derived from the self-compatible, yellow-seeded subspecies *trilocularis* (Stephenson et al. 2010). Because the TILLING population is heavily mutagenized, putative loss-of-function alleles were backcrossed six times to the parental line before generating homozygous mutants. For all crosses, flowers were emasculated prior to anthesis and were pollinated one day later. The resulting plants were genotyped with molecular markers using standard approaches. *braA.nrpd1.a-2* (at BraA09006512) was genotyped with AAACGGGAGACAGCTTCTTACG and AGGAACGTACCCGTGAAGACAGACT and digested with HinfI; *braA.rdr2.a-2* (at BraA09004909) was amplified with TTCGAAATGTGCTGCTAGGATGAGC and CTGAAGGAACTGCGGTCAACG and digested with AluI; *braA.nrpe1.a-1* (at BraA03002228) was amplified with CTGGCCCGGCTTGGACATTTTTCC and ACAAGTCTTCCAGACTTAAACTAAATCCTT and digested with AflII.

To assess embryogenesis, seeds were manually dissected at 17 days after fertilization, or were cleared as described in (Wang et al.). *B. rapa* and Arabidopsis seed weight was measured from seed that had dried for at least four weeks. *B. rapa* seeds were measuring in pools of 5, while Arabidopsis seeds were measured in pools of 10.

### Small RNA sequencing and analysis

Unfertilized ovules from *braA.nrpd1.a-2*, *braA.rdr2.a-2*, and *braA.nrpe1.a-1* were collected less than 24 hours prior to anthesis and total RNA was prepared in a method adapted from (White and Kaper 1989). Briefly, frozen tissue was ground to a fine powder and mixed with at least 5 volumes of extraction buffer (100 mM Glycine, 10 mM EDTA, 100 mM NaCl, 2% SDS, pH 9.5), prior to organic extraction and precipitation. Small RNA was enriched with a mirVana miRNA isolation kit (Thermo Fisher Scientific) prior to NEBNext small RNA library preparation (New England Biolabs). Three independent biological replicates were prepared for each genotype. Barcoded libraries were pooled and sequenced on an Illumina HiSeq 2500 at The University of Missouri DNA Core Facility.

Trimmed reads from unfertilized ovules were aligned to the *B. rapa* R-o-18 genome (version 1.2, A Baten and GJ King, unpublished) using Bowtie (Langmead et al. 2009). Only alignments with no mismatches were allowed. Genome-mapping reads were further filtered by aligning to the Rfam database v12.1 (Nawrocki et al. 2015) and removing small RNAs matching structural and noncoding RNAs (excluding miRNAs and miRNA precursors) annotated in *B. rapa* and *A. thaliana*. Reads mapping to the *A. thaliana* chloroplast and mitochondrial genomes (TAIR 10), and reads <19 or >26 nt were also removed (Supplementary Table 2). In all cases, no mismatches were allowed in alignments. Size profiles of independent replicates were similar, and therefore replicates were pooled for further analyses.

For small RNA locus annotation, filtered reads from pooled wild-type libraries were mapped to the R-o-18 genome using ShortStack v3.8.1 (Axtell 2013; Johnson et al. 2016) allowing no mismatches. A minimum read depth of 2 reads per million (approximately 12 reads in wild type) was used to define sRNA loci. Pooled libraries were then realigned with ShortStack to compare expression at wild-type-defined loci. To address over-sampling, raw read counts were normalized by the number of mapped 21 nt reads in each pooled library.

Reads from intraspecific crosses were filtered and mapped as above except both the R-o-18 and R-500 genomes (K Greenham and CR McClung, unpublished) were used for mapping (Supplemental Table 3). Genome-specific reads were defined as reads with a perfect match in only one of the two parental genomes.

Small RNA sequence data were deposited at the NCBI Short Read Archive (SRA) under accession numbers SRP114437 (RdDM mutants) and SRP114469 (intraspecific crosses).

## Acknowledgements

We are grateful to Dr. Robertson McClung and Dr. Kathleen Greenham from Dartmouth University for sharing the *B. rapa* R-500 genome prior to publication. Thanks also to Fran Robson at RevGenUK for TILLING advice. This research was funded by the NSF Plant Genome Research Program (IOS-1546825 to R.A.M).

## List of supplemental materials

Supplemental Table 1. *B. rapa* RdDM mutant alleles.

Supplemental Table 2. Small RNA sequencing of RdDM mutants.

Supplemental Table 3. Small RNA sequencing of reciprocal crosses.

